# Characterization of a novel variant in the HR1 domain of *MFN2* in a patient with ataxia, optic atrophy and sensorineural hearing loss

**DOI:** 10.1101/2021.01.11.426268

**Authors:** Govinda Sharma, Rasha Saubouny, Matthew M Joel, Kristina Martens, Davide Martino, A.P. Jason de Koning, Gerald Pfeffer, Timothy E. Shutt

**Affiliations:** Departments of Medical Genetics and Biochemistry & Molecular Biology, Cumming School of Medicine, Alberta Children’s Hospital Research Institute, Hotchkiss Brain Institute, University of Calgary; Departments of Clinical Neurosciences and Medical Genetics, Cumming School of Medicine, University of Calgary, Hotchkiss Brain Institute, Alberta Child Health Research Institute; Department of Biochemistry & Molecular Biology, Cumming School of Medicine, Alberta Children’s Hospital Research Institute, University of Calgary; Department of Clinical Neurosciences, Cumming School of Medicine, Hotchkiss Brain Institute, University of Calgary

## Abstract

Pathogenic variants in *MFN2* cause Charcot-Marie-Tooth disease (CMT) type 2A (CMT2A) and are the leading cause of the axonal subtypes of CMT. CMT2A is characterized by predominantly distal motor weakness and muscle atrophy, with highly variable severity and onset age. Notably, some *MFN2* variants can also lead to other phenotypes such as optic atrophy, hearing loss and lipodystrophy. Despite the clear link between *MFN2* and CMT2A, our mechanistic understanding of how dysfunction of the MFN2 protein causes human disease pathologies remains incomplete. This lack of understanding is due in part to the multiple cellular roles of MFN2. Though initially characterized for its role in mediating mitochondrial fusion, MFN2 also plays important roles in mediating interactions between mitochondria and other organelles, such as the endoplasmic reticulum and lipid droplets. Additionally, MFN2 is also important for mitochondrial transport, mitochondrial autophagy, and has even been implicated in lipid transfer. Though over 100 pathogenic *MFN2* variants have been described to date, only a few have been characterized functionally, and even then, often only for one or two functions. Here, we describe a novel homozygous MFN2 variant, D414V, in a patient presenting with cerebellar ataxia, deafness, blindness, and diffuse cerebral and cerebellar atrophy. Characterization of patient fibroblasts reveals phenotypes consistent with impaired MFN2 functions and expands the phenotypic presentation of *MFN2* variants to include cerebellar ataxia.

## Introduction

Mitochondria are highly dynamic double membrane-bound organelles that undergo continuous remodelling via fusion and fission events. These dynamic processes determine mitochondrial structure and regulate mitochondrial function (1). Mitochondrial fusion is a multistep process mediated by several essential proteins and regulators (2). Tethering of adjacent mitochondria and fusion of outer mitochondrial membrane (OMM) is performed by mitofusin1 and mitofusin2 (MFN1/2), two homologous proteins with partially redundant functions that are integral to the OMM. Meanwhile, fusion of the inner mitochondrial membrane (IMM) is carried out by optic atrophy 1 (OPA1) (3, 4). Highlighting the importance of mitochondrial fusion is the fact that knockout of *MFN1*, *MFN2* or *OPA1* genes is embryonic lethal (5, 6). In addition, pathogenic variants in these genes cause human disease, with *OPA1* linked to optic atrophy, and *MFN2* linked to the peripheral neuropathy Charcot Marie Tooth type 2A (CMT2A) (7, 8).

MFNs have an N-terminal GTPase domain that is exposed to the cytosol, and two heptad repeat domains (HR1 and HR2, also referred to as coiled-coil domains), separated by a transmembrane domain. While several structural models have been generated for MFNs (9–12), these are from artificial constructs that are based on the notion that N-terminal and C-terminal domains of the protein both face the cytosol and can interact. However, a recent reappraisal of the topology of MFNs showed that the C-terminus of the protein, including the HR2 domain, is exposed to the inner membrane space (IMS), not the cytosol (13). This revision raises questions about the validity of the structural models. Thus, there remain many questions as to the exact structure of MFNs and how they mediate mitochondrial fusion.

Mitochondrial fusion is also important for the maintenance of the mitochondrial genome (mtDNA), which is present in hundreds to thousands of copies per cell. The mtDNA is packaged into nucleoid structures, each of which is approximately 100 nm in size and contains a single copy of the mitochondrial genome (14–16). Impairments to mitochondrial fusion can lead to reduced copy number and increased nucleoid size (17, 18). While the reduced copy number is thought to be a result of decreased distribution of the replication machinery (19), it is unclear exactly why nucleoid size might change. Meanwhile, mitochondrial respiration can also be compromised upon loss of fusion (20, 21).

Notably, MFN2 has several functions in addition to its role in mitochondrial fusion. For example, MFN2 mediates mitochondrial autophagy (22, 23) and transport of mitochondria (24, 25). Meanwhile, MFN2 also localizes to the endoplasmic reticulum, where it mediates interactions between the ER and mitochondria (26–28). These mitochondria-ER contact sites (MERCs) are specialized sites for both lipid biogenesis and exchange, are important for regulating calcium signalling, and can also mark sites of mitochondrial fission (29), as well as mtDNA replication (30). Notably, MFN2 has been implicated in directly binding and transferring phosphatidylserine from ER to mitochondria (31). While there is some debate whether MFN2 promotes or inhibits MERCs (27, 28, 32, 33), it is clear that it plays an important role in these organelle contacts. Finally, in addition to MERCs, MFN2 also mediates interactions between mitochondria and lipid droplets (34).

To date, more than 100 pathogenic variants in *MFN2* have been associated with peripheral neuropathy (35). However, despite the general assumption that impaired mitochondrial fusion causes the peripheral neuropathy phenotype, only a few pathogenic MFN2 variants have been investigated functionally for their effects on MFN2 functions. While some pathogenic MFN2 variants do impair fusion (36, 37), unexpectedly, other pathogenic variants seem to increase fusion (38, 39), while several pathogenic variants do not appear to affect fusion at all (37, 40, 41). These findings raise the possibility that impaired fusion does not lead to peripheral neuropathy *per se*.

In this context, it is notable that other MFN2-mediated mitochondrial functions can also be impacted by pathogenic MFN2 variants. For example, disruptions to MERCs (26, 37), lipid metabolism (42), mitochondrial respiration (43), mtDNA copy number (44–46), and mitochondrial transport (25), have all been observed in association with various pathogenic MFN2 variants. However, it should be noted these phenotypes have not all been widely investigated across a variety of MFN2 variants.

Further complicating our mechanistic understanding of how MFN2 dysfunction causes disease is the fact that additional pathogenic phenotypes can also be linked to MFN2 variants. Although not common, some MFN2 variants are linked to other disease phenotypes such as optic atrophy (47, 48). Central nervous system involvement is also rarely described (49), with periventricular and subcortical white matter lesion pathogenic for age, associated with transient neurological deficits in a small minority of cases. Other complex phenotypes have been observed in the presence of homozygous or compound heterozygous mutations. For example, the R707W *MFN2* variant, which causes CMT2A when heterozygous, is also associated with lipodystrophic syndromes when homozygous. Specifically, R707W causes proliferation of adipocytes leading to adipose hyperplasia and lipomatosis (42, 50, 51). Additional atypical features including severe neuropathy with hearing loss has been described with biallelic mutations (47).

In summary, MFN2 performs a number of functions, many of which can be impaired by pathogenic variants in *MFN2*. Meanwhile, *MFN2* variants can lead to a number of different patient phenotypes. However, there is no clear understanding of the molecular mechanisms causing disease and whether impairment of specific MFN2 functions leads to specific phenotypes. Here, in a patient presenting with ataxia, sensorineural hearing loss, and optic atrophy leading to vision loss, we report the presence of a homozygous novel candidate pathogenic variant in MFN2, p.(D414V), located in the middle of the HR1 domain. While ataxia has not been associated previously with MFN2 specifically, it is common in mitochondrial disease. Our characterization of mitochondrial functions in patient fibroblasts is consistent with impairment of MFN2 functions, expanding the clinical spectrum of phenotypes associated with pathogenic variants in *MFN2*.

## Results

### Ethics statement

This study was approved by the University of Calgary Conjoint Health Research Ethics Board with the following approval numbers: REB15-2763 (for exome sequencing), REB17-0850 (for the skin biopsy/fibroblast lines). Written informed consent was obtained from the participant for both of the above projects.

### Case description and clinical diagnosis

The patient initially came to medical attention at eight years of age, having previously had a normal birth history and development. He began to experience visual blurring affecting both eyes, and bilateral optic atrophy was identified on ophthalmologic evaluation. His vision continued to slowly deteriorate over time. In his 20’s he developed gait disturbance which was diagnosed as ataxia (though in retrospect teachers had informed him of changes to his gait as early as 12 years of age). The ataxia progressed to affect limb coordination functions in his 30’s, in association with dysarthria. Sensorineural hearing loss was diagnosed in his 20’s and progressed gradually to deafness. In his 40’s he developed hypertension as well as type 2 diabetes mellitus. He was coincidentally diagnosed with a painful small-fibre sensory neuropathy, which was thought to be attributable to his diabetes.

His family history did not include any other individuals with neurologic disease. His parents originated from India and no consanguinity was reported. His two siblings are healthy and do not have a similar condition. The patient does not have any children.

By the time of his assessment by one of the authors (GP) at age 54 years, his family described severe visual and hearing loss. He was bound to his wheelchair and required assistance for feeding and all daily living tasks due to severe balance and coordination deficits. On neurologic examination, he was alert and attentive. His cognition was difficult to assess due to the above-mentioned visual/hearing loss and communication difficulties, but his overall cognition seemed grossly appropriate. Cranial nerve examination demonstrated bilateral optic atrophy, light perception only for visual acuity, full extraocular movements, no ptosis, and symmetric facial movements. His hearing was poor, but he was able to comprehend speech when spoken in a loud voice directly into his ear. He also had cerebellar dysarthria, but speech was still intelligible. He otherwise had normal lower cranial nerve functions. He had diffuse and symmetric loss of muscle bulk which was attributed to deconditioning, with normal motor tone. Deep tendon reflexes were unobtainable in all extremities. Plantar responses were flexor. Power examination revealed that muscle strength was well preserved, with some minor hip girdle weakness which was again attributed to disuse. Additionally, he exhibited a postural tremor in his upper limbs and bilateral dysmetria and intention tremor on finger-to-nose testing. He required two-person assistance to stand. Sensory examination was fairly unremarkable, with vibration sensory loss below the ankles bilaterally, but was otherwise normal.

The differential diagnosis at this time was considered to most likely include genetic or metabolic disorders, such as mitochondrial disorders, CAPOS syndrome (cerebellar ataxia, areflexia, pes cavus, optic atrophy, and sensorineural hearing loss, due to mutations in *ATP1A3*), or other multisystem genetic disorders such as Wolfram syndrome (due to *WFS1* mutations). Investigations included vitamin B12, folate, and vitamin E levels; which were normal. Very long chain fatty acids and hexosaminidase A and B were also normal. Testing for anti-GAD antibodies was negative. A muscle biopsy demonstrated some denervation atrophy, but did not reveal any changes suggesting a mitochondrial cytopathy. MRI of the brain revealed diffuse cerebral volume loss affecting the cerebrum, cerebellum and brainstem structures, and volume loss was also observable in the optic nerves bilaterally and optic chiasm (Fig. 1A). Clinical genetic testing included a first-line genetic screening for spinocerebellar ataxia types 1, 2, 3, 6, 7, 8, and Friedreich ataxia, which was negative. He also had an NGS-based sequencing panel for ataxic syndromes including 277 genes. This found a homozygous variant of unknown significance in *MFN2*, c.1241A>T p.(D414V).

**Figure 1.**
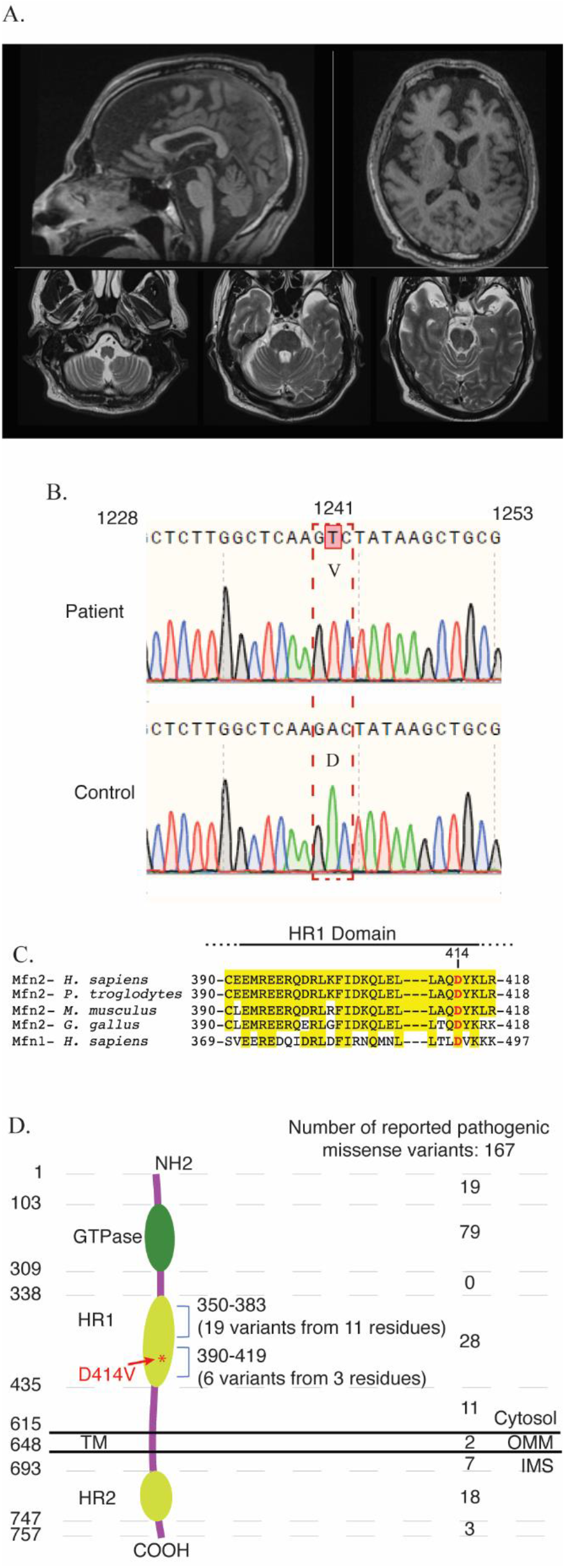
Pathogenic variants in *MFN2*. Representative images of MRI head scans of the patient at 55 years of age. Top-left: T1-weighted sagittal image through the midline demonstrates volume loss in the frontal lobe, brainstem, and midline cerebellar structures. Top-right: T1-weighted transverse axial image at the level of the lateral ventricles and basal ganglia demonstrate volume loss predominantly affecting the bilateral frontal lobes. Bottom panels: T2-weighted transverse axial images at the levels of the medulla and pons again demonstrate brainstem atrophy as well as accentuated cerebellar foliae indicative of diffuse volume loss in the posterior fossa. (B) Sequencing chromatograms confirms the 1241A>T variant, resulting in a missense mutation of Aspartic acid (D) to Valine (V) at position 414 in the MFN2 protein. Sequencing data are shown from patient derived (upper panel) or normal control (lower panel) derived fibroblasts. (C) Alignment of the region of the HR1 domain of MFN2, showing D414V is conserved throughout vertebrate species and with MFN1. Residues highlighted in yellow are conserved residues determined by Clustal Omega analysis. (D) Diagram showing the topology and domains of the 757 amino acid MFN2 protein, which contains a GTPase domain, two HR domains and a transmembrane (TM) domain. The number of reported pathogenic missense mutations in the indicated regions of the protein are indicated on the right.

Exome sequencing was performed and variants were filtered as follows: maximum allele frequency in population databases of <0.0001, predicted to cause protein-coding changes (simple substitutions, frameshifts, splicing alterations, or early termination), and present in genes associated with neurological phenotypes. We considered variants to be reasonable candidates if they met these criteria and if they fit the known mode of inheritance for these conditions (e.g: recessive disease genes would require two heterozygous variants or a homozygous variant), and were not classified as “benign” or “likely benign” in ClinVar. Using these criteria, we again identified the above-mentioned homozygous variant in *MFN2* (c.1241A>T, p.(D414V)). Using these criteria, we did not identify any other monogenic disease candidates.

### Characterization of patient fibroblasts

We obtained skin fibroblast cells from the patient in order to examine whether the D414V variant affects the various functions of MFN2 and is thus likely to be pathogenic. We first confirmed the presence of MFN2-D414V variant in the patient derived fibroblasts by Sanger sequencing (Fig. 1B). The aspartate residue at 414 position of MFN2 is conserved throughout vertebrate orthologs, as well as the human MFN1 paralog (Fig. 1C). This variant falls in the HR1 domain of the MFN2 protein (Fig. 1D). Notably, a previous bioinformatic study predicted that this amino acid change would be damaging (52).

### Impact of the MFN2-D414V variant on mitochondrial dynamics

We compared mitochondrial morphology in patient and control fibroblasts in order to examine the well-defined role for MFN2 in mediating mitochondrial fusion. While the mitochondria were long and often reticular in the control fibroblasts, those in the patient fibroblasts were noticeably shorter (Fig 2A). We quantified this observation by measuring the mitochondrial length and found shorter mitochondria in patient fibroblasts, indicative of more fragmented mitochondrial networks compared to those of control fibroblasts (Fig. 2B). These results are consistent with a reduction in mitochondrial fusion.

**Figure 2.**
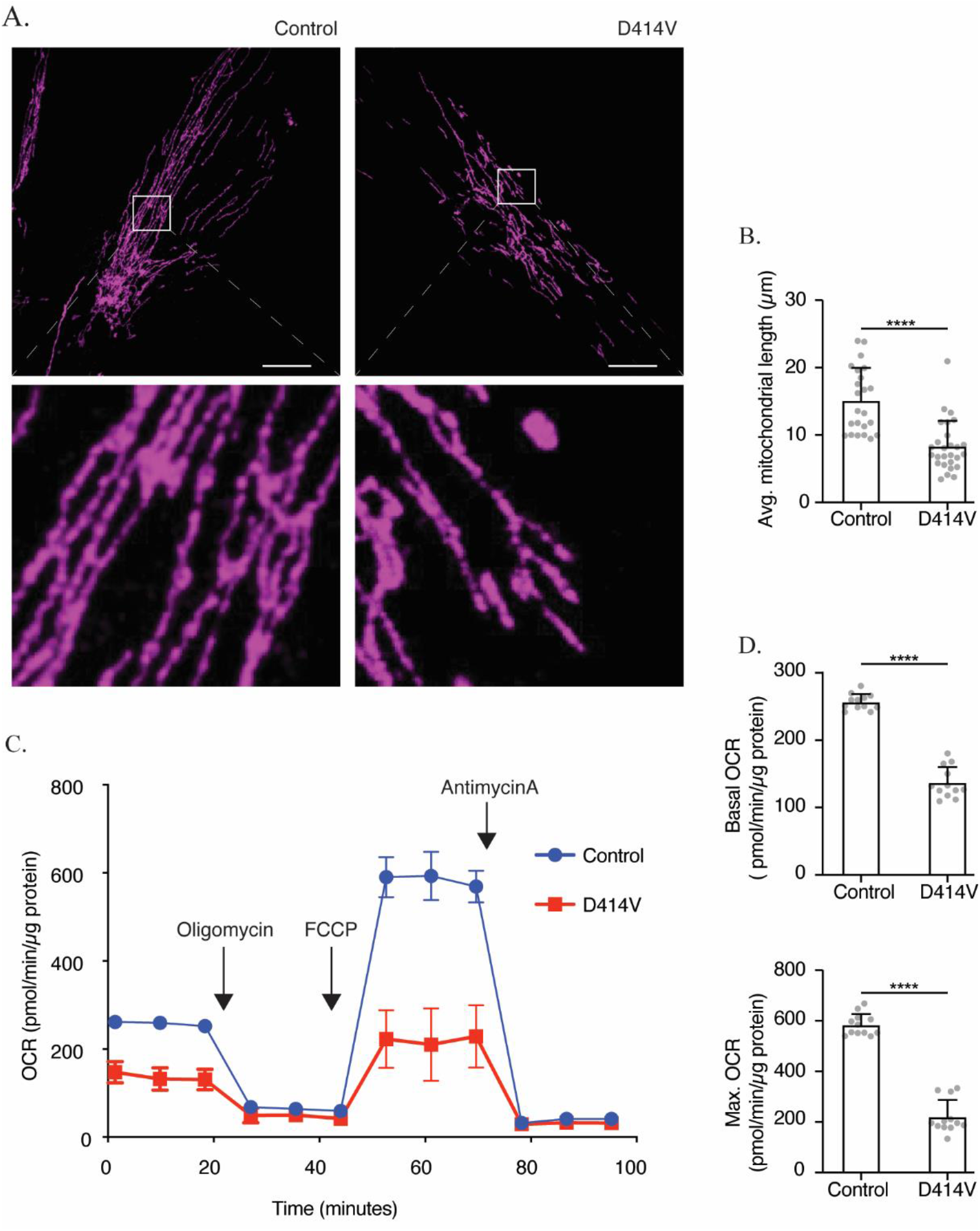
Mitochondrial dysfunction in *MFN2*-D414V patient-derived fibroblasts. (A) Representative confocal images of healthy control (left panel) and patient derived (right panel) fibroblasts labelled with TOMM20 immunostaining (purple) to visualize mitochondrial networks. Scale bar, 20 μm. (B) Quantification of average mitochondrial length in control and patient cells. The average mitochondrial lengths of 24 control and 27 patient fibroblasts. Error bars indicate mean +/− SD. p<0.0001, unpaired t-test (C) Oxygen consumption rate (OCR) traces in control and MFN2-D414V fibroblasts measured using the Seahorse XF24 extracellular flux analyzer. (D) Basal (upper) and maximum (lower) OCR in control and Mfn2-D414V fibroblasts calculated from C. Error bars indicate mean +/− SD. p<0.0001, unpaired t-test.

### Mitochondrial oxygen consumption rate in patient fibroblasts

Given the links between mitochondrial form and function, we checked whether the D414V MFN2 variant might impact mitochondrial bioenergetics by measuring the oxygen consumption rate. Notably, both basal and maximal oxygen consumption rates were significantly lower in patient fibroblasts when compared to control (Fig 2C, D).

### Mitochondrial nucleoids

In order to understand how the D414V variant might be impacting mitochondrial respiration, we employed several approaches to examine the mtDNA genome, which is essential for respiration, and which can be impacted by impairments to mitochondrial fusion (19). First, using confocal microscopy, we observed a significant decrease in the average number of nucleoids in patient fibroblasts compared to control (Fig. 3A, B). Given previous findings that impaired mitochondrial fusion can lead to larger mitochondrial nucleoids, we also imaged and quantified the size of mtDNA nucleoids. Unexpectedly, we observed a reduction in the average size of nucleoids in patient fibroblasts when compared to those from control (Fig. 3C). Consistent with the reduced number of nucleoids evident by confocal microscopy, quantitative PCR analysis also showed a slight reduction in mtDNA copy number in patient fibroblasts (Fig. 3D). Meanwhile, D414V patient fibroblasts did not exhibit any evidence for mtDNA deletions (Fig 3E), as has been reported previously for some pathogenic MFN2 variants (48).

**Figure 3.**
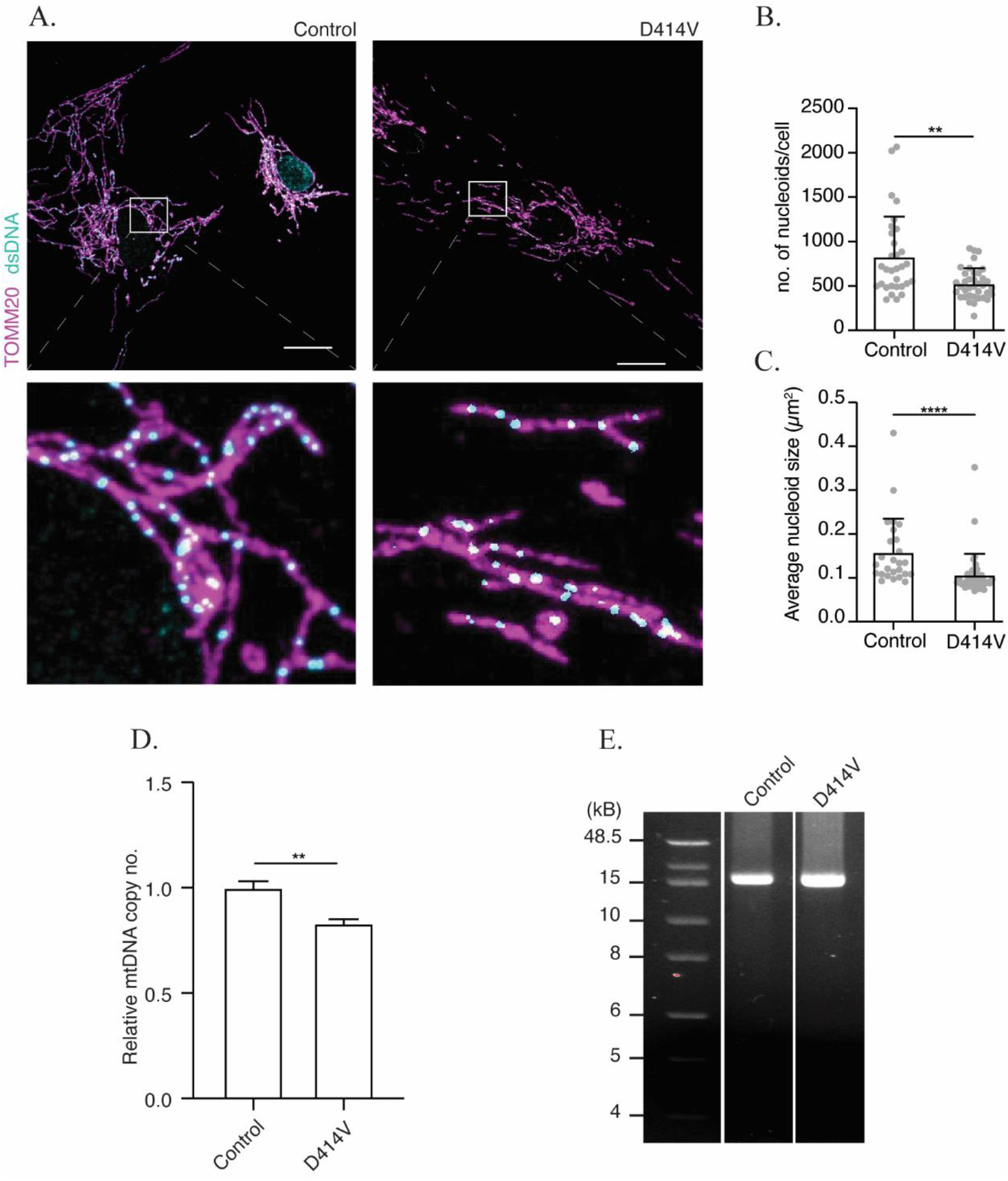
Altered mtDNA in MFN2 D414V fibroblasts. (A) Representative confocal images of mitochondria and mitochondrial nucleoids labelled by immunostaining with anti-TOMM20 (purple) and anti-dsDNA (cyan) antibodies in control and *MFN2*-D414V fibroblasts. The area indicated by white box in the upper panel is shown in lower panel at higher magnification. Scale bar, 20 μm. (B) Quantification of the number of mtDNA nucleoids. Each data point indicates the total number of nucleoids in a control or MFN2 D414V fibroblast. Error bars indicate mean +/− SD. p=0.0012, unpaired t-test. (C) Quantification of the average mtDNA nucleoid size in control (n, 26) or MFN2-D414V fibroblast cells (n, 40). Error bars indicate, mean +/− SD. p<0.0001, unpaired t-test. (D) Relative mtDNA copy number analyzed using quantitative PCR. (E) Integrity of the mtDNA genome was assessed on 0.6% agarose gel following long-range PCR of DNA isolated from control or MFN2-D414V fibroblasts. No mtDNA deletions were detected.

### Mitochondria-ER contacts (MERCs)

Next, we tested whether the D414V variant might impair MFN2 protein function in mediating MERCs. To estimate the size and number of MERCs in patient and control fibroblasts, we employed a proximity ligation assay (PLA), which indicates when two target proteins (mitochondrial-TOMM20 and ER-calnexin) are within ~40nm, (53, 54). We observed a significant reduction in both the number and size of MERCs in the patient cells (Fig 4A-D), consistent with the notion that the D414V variant might be causing reduced interactions between mitochondria and ER.

**Figure 4.**
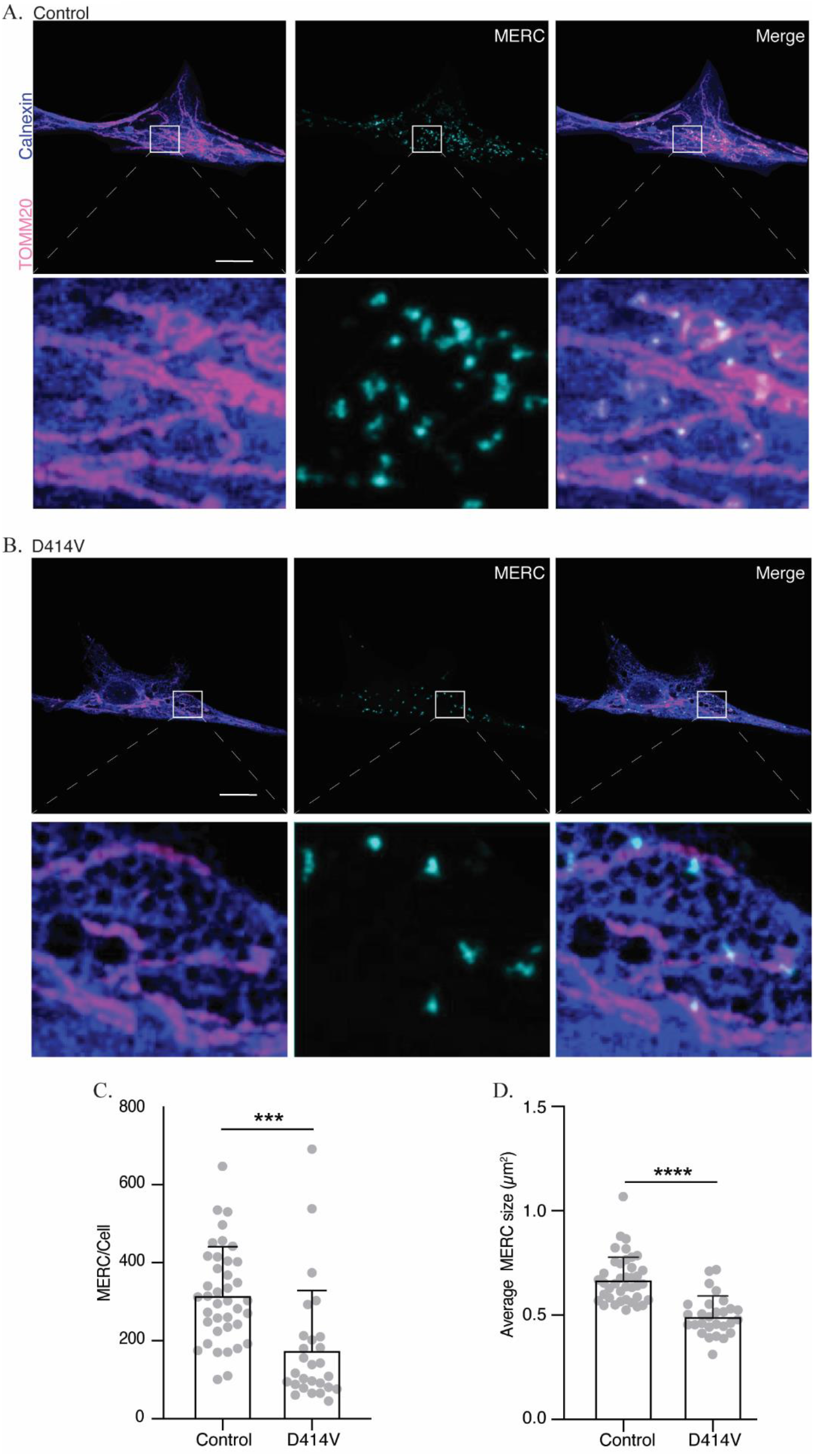
Mitochondria-ER contacts (MERCs) are reduces in MFN2 D414V fibroblasts. (A and B) Representative images showing MERCs in the control (A) or MFN2 D414V (B) fibroblasts, visualized using a proximity-ligation assay (cyan). Mitochondrial networks and ER are visualized via immunostaining with TOMM20 (pink) and calnexin (purple), respectively. The area indicated by white box in the upper panel is shown in lower panel zoomed in. Scale bar, 20 μm. (C) Quantification of the number of MERCs. Each data point indicates the total number of MERCs from an individual control or MFN2-D414V fibroblast cell. Error bars indicate SD. p=0.0002, unpaired t-test. (D) Quantification of the average size of MERC’s in control or MFN2-D414V fibroblasts. Each data point indicates average size of MERCs from an individual cell. Error bars indicate SD. p<0.0001, unpaired t-test.

### Lipid droplet regulation

Our observations of reduced MERCs, and the probable compromise in tethering function of MFN2 in the patient fibroblasts, led us to hypothesize that lipid droplet metabolism could also be affected by the MFN2-D414V variant. To this end, lipid droplets were imaged by staining with a neutral lipid dye and imaged by confocal microscopy. Notably, the number of lipid droplets was reduced in the patient fibroblasts (Fig. 5A, B and C). In addition, the total lipid content, as measured by the total intensity of the lipid dye, was also reduced (Fig. 5D). Finally, we also noted an unusual perinuclear arrangement of lipid droplets in patient fibroblasts compared to control fibroblasts, in which lipid droplets were spread throughout the cell (Fig. 5A, B). To quantify the differences in lipid droplet distribution, we measured the distance of individual lipid droplets from the center of the nucleus. We found that the average distances of lipid droplets from the nucleus was significantly lower in the patient fibroblasts (Fig. 5E, F). Altogether, these observations are consistent with the notion that the D414V variant in MFN2 affects lipid droplets.

**Figure 5.**
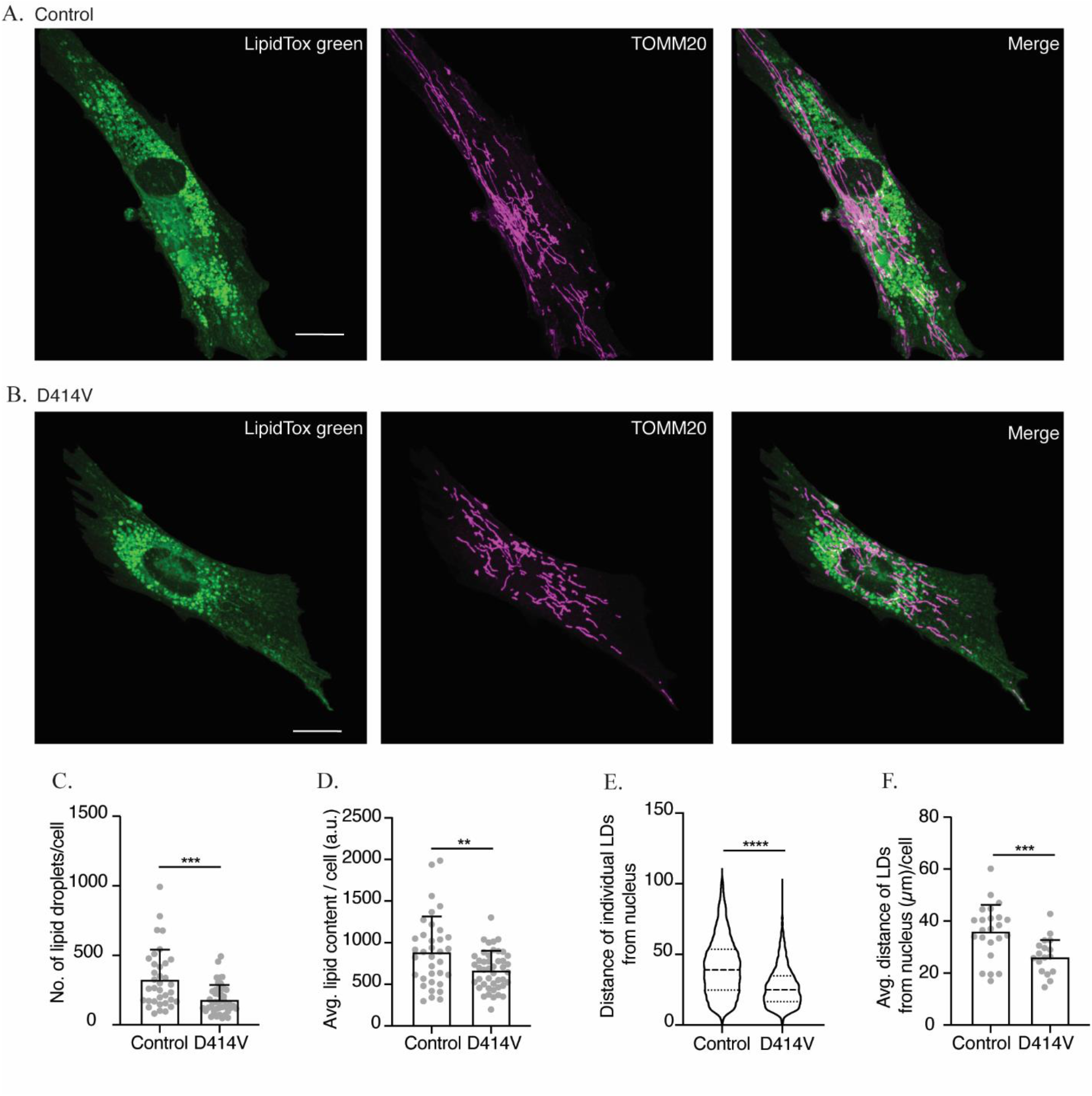
The intracellular distribution and number of lipid droplets (LD) are reduced in MFN2 D414V fibroblasts. (A and B) Representative confocal images of control (A) and MFN2 D414V (B) fibroblasts, with lipid droplets stained with LipidTox Green (green) and mitochondria visualized following immunofluorescence with TOMM20 (purple). The area indicated by white box in the upper panel is shown in lower panel zoomed in. Scale bar, 20 μm. (C) Quantification of the number of LDs. Each data point indicates the total number of LDs in a control or MFN2 D414V fibroblasts. Error bars indicate SD. p=0.0002, unpaired t-test. (D) Quantification of total lipid content. Each data point indicates the total intensity given by the LipidTox Green dye from each of the control or *MFN2* D414V fibroblasts. Error bar indicates SD. p=0.0073, unpaired t-test. (E) Quantification of the intracellular distribution of LDs. Distance of each LD from centre of the nucleus of a respective cell was calculated for control (n, 8303) and MFN2-D414V (n, 5259). The median and inter-quartile ranges are indicated in the violin plot. p<0.0001, unpaired t-test. (F) The average distance of LDs from nucleus. Each data point indicates the average distance of all the LDs from the center of respective cell’s nucleus for control or MFN2 D414V. Error bar indicates SD. p=0.0006, unpaired t-test.

## Discussion

We identified a novel homozygous *MFN2* variant, D414V, in a patient with ataxia, deafness and blindness. While hearing and vision loss have been observed in a few cases linked to *MFN2* variants (47, 48), cerebellar ataxia has not. As pathogenic variants in MFN2 are typically associated with the peripheral neuropathy CMT2A, it was not clear whether the D414V variant is responsible for these patient phenotypes. Notably, though the patient did have a small-fibre sensory neuropathy, this was attributed to diabetes given the relatively late onset compared to the other phenotypes. A second question arising from the D414V variant is whether different mechanisms underlying MFN2 dysfunction might explain the distinct patient phenotypes. Here, we discuss our characterization of the D414V variant, and the novel insight provided into the function of MFN2 and molecular mechanisms underlying pathology associated with dysfunction of this protein.

Despite the limitations of existing structural models, homology modelling of MFN2 with the bacterial homolog BDLP may still be informative (55, 56). These reports suggest that the HR1 domain, which is predicted to form part of a larger helical bundle, comprises two alpha helical subdomains (~350-383, and ~390-419) that are separated by a flexible hinge domain. Notably, of the 167 reported MFN2 missense/nonsense variants in the Human Gene Mutation Database (57), only a handful are found in at only three positions in the second helical domain of HR1 (aa390-419), with two of the affected residues at the extremities of this subdomain (i.e. aa390 and aa 418). These variants have been published in the context of classic CMT2A phenotypes, including: C390R (47), C390F (8), R400X (58), R400P (59), R418Q (60) and R418X (61). However, none of these variants have been characterized functionally in human cells. Moreover, in contrast to the homozygous D414V reported here, the other published variants in the second helical domain of HR1 are all heterozygous. Notably, two variants were compound heterozygous, with the R400X variant found in combination with another pathogenic variant (R250W, also linked to CMT2A), and the C390R variant in combination with N214D. As such, some functional rescue by the other allele is possible for these other variants. These observations suggest that the region comprising the second helical domain within HR1 (aa390-419), where the D414V variant is present, is of critical importance for proper MFN2 function.

It is also worth briefly discussing a study in which two rare human *MFN2* variants were modelled in Drosophila (52). Tissue-specific overexpression of these variants (M393I and R400Q) led to cardiac and eye phenotypes. Notably, these variants are also present in the second helical region of the HR1 domain, suggesting a distinct role for this region of the HR1 domain in mediating MFN2 function, and hint at additional phenotypes that could be associated with human disease due to pathogenic variants in *MFN2*.

To investigate whether the D414V variant is indeed pathogenic, we performed a comprehensive analysis of mitochondrial functions linked to MFN2, beginning with the well-recognized role of MFN2 in mediating mitochondrial fusion. Patient fibroblasts harboring the D414V variant displayed fragmented mitochondrial networks compared to control, consistent with impaired fusion in these cells. This finding is notable in comparison to the few pathogenic MFN2 variants causing CMT2A that have been investigated for their role in fusion. While a few of the CMT2A *MFN2* variants studied appear to be fusion incompetent, (5, 52, 62), there are other CMT2A *MFN2* variants that do not seem to affect mitochondrial morphology (40, 63). Conversely, some CMT2A MFN2 variants actually enhance fusion (38, 39). Thus, impaired mitochondrial fusion does not appear to be necessary to cause CMT2A.

As mitochondrial fusion is important for maintenance of the mitochondrial genome, we also investigated mtDNA in patient fibroblasts. The reduced mtDNA copy number and nucleoid number observed in patient fibroblasts is consistent with impaired mitochondrial fusion in these cells, as reduced copy number is proposed to be due to inefficient fusion-dependent distribution of the mtDNA replication machinery (19). Surprisingly, we also observed slightly smaller nucleoids in patient cells, an unexpected finding given that impaired mitochondrial fusion has previously been linked to enlarged mitochondrial nucleoids (19, 21, 64). However, it should be noted that it is not clear why nucleoid size increases in response impaired fusion.

One possible explanation for this discrepancy in nucleoid size could be that size does not correlate directly with the degree of fusion impairment. Perhaps, slight impairments to mitochondrial fusion lead to smaller nucleoids, while greater impairments lead to larger nucleoids. The fact that each nucleoid is estimated to contain ~1.4 copies of the mtDNA genome (65), is consistent with observations that a significant subset of nucleoids is actively undergoing replication (30). If one logically assumes that replicating nucleoids are larger than non-replicating nucleoids, then the smaller nucleoids we observe could be due to a reduced number of nucleoids being actively replicated.

Meanwhile, the larger nucleoids described in cells completely lacking fusion are due clustering of multiple individual nucleoids that cannot be resolved by traditional confocal microscopy (Silva Ramos et al., 2019). Notably, cells where mitochondrial fission is inhibited also exhibit large clustered nucleoids, demonstrating that mitochondrial dynamics are important for the distribution of nucleoids (66, 67). Perhaps in cells with more severe fusion impairment, individual mtDNA nucleoids also cluster, leading to apparently larger nucleoids. It is also notable that while nucleoid size has not previously been quantified in cells with different pathogenic MFN2 variants, a subset of pathogenic MFN2 variants are linked to mtDNA depletion and deletions (46, 68).

Given the links between mitochondrial structure and function, as well as the critical role for mtDNA encoded proteins in oxidative phosphorylation, we also examined mitochondrial function in patient fibroblasts. The reduced OCR we observed in *MFN2*-D414V patient fibroblast demonstrated clear mitochondrial dysfunction in these cells, possibly as a result of reduced mitochondrial fusion leading to reduced levels of mtDNA, or impaired MERCs affecting Ca^++^ transfer to mitochondria. While MFN2 is important for maintaining mitochondrial bioenergetics (69), the consequences of pathogenic CMT2A *MFN2* variants on mitochondrial function are conflicting, raising questions about the contribution of impaired mitochondrial bioenergetics to CMT2A. For example, some pathogenic CMT2A *MFN2* variants lead to a decrease in the mitochondrial bioenergetic function (40, 69, 70), others cause an increase (71, 72), while still more variants do not cause any change at all (25, 37, 41).

MFN2 also plays a key role in mediating contact sites between mitochondria and ER, though conflicting reports debate the exact role of MFN2 in maintaining MERCs. Thus, we also investigated MERCs in D414V patient fibroblasts, where we observed a decrease in number as well as size. These results suggest that the D414V variant reduces MERCs. While only a limited number of CMT2A MFN2 variants have been investigated for their role in MERCs (26, 37), disruption of contacts between mitochondria and ER seems to be a common theme underlying neuropathies (62, 73, 74). Further, a study in *Drosophila* indicated that MFN2 could regulate ER stress (75), and peripheral neuropathy in diabetic patients is also thought to be partly due to ER stress (76, 77). These observation suggest that deregulation of MERCs and their many functions could be a significant contributor to the peripheral neuropathy phenotype associated with CMT2A *MFN2* variants (37, 62).

Although a peripheral neuropathy noted in the D414V patient was initially attributed to a diabetic neuropathy, it is worth considering the contribution of MFN2 dysfunction to this phenotype. Notably, peripheral neuropathy can appear at later stages in life and with varying severity (35, 78). Phenotypic variation may occur between individuals with the same mutation suggesting a potential role for other genetic or environmental factors. In this regard, the alteration to MERCs are consistent with the fact that impaired MERCs are a feature of MFN2 CMT2A variants (37, 79). As such, it may simply be a coincidence that the peripheral neuropathy appeared around the same time as the patient’s diabetes, and that MFN2 dysfunction was responsible. Alternatively, it is possible that the development of diabetes exacerbated an already unstable situation due to the *MFN2* variant.

Next, we examined lipid droplets, another organelle whose interactions with mitochondria are mediated by MFN2 (34). We found that the abundance of neutral lipid signal and the number of lipid droplets are reduced in patient fibroblasts. In contrast to our finding, a previous report has shown that CMT2A *MFN2* variants increase the lipid droplet signal and apparent abundance (37). Interestingly, we also discovered an unexpected perinuclear accumulation of lipid droplets in fibroblasts with the D414V variant when compared to healthy control. This phenotype has not been described previously in the context of MFN2 dysfunction.

It is also worth discussing the role of MFN2 in cellular lipid homeostasis, though there is clearly more to learn, especially in the context of disease. In addition to regulating lipid droplets and MERCs, MFN2 was recently shown to have a direct role in transferring phosphatidylserine from the ER to mitochondria, a function implicated in non-alcoholic steatohepatitis (31). Moreover, in adipose tissue the association between mitochondria and lipid droplets is important for both lipid storage and consumption (80, 81), while adipocyte specific knockout of *MFN2* leads to obesity in mice (34, 82, 83). Finally, the R707W MFN2 variant, which causes CMT2A when present heterozygously, also causes lipomatosis when present homozygously (50, 51). Thus, it is clear that the roles of MFN2 in maintaining lipid homeostasis are important, though the exact molecular mechanisms remain undefined.

Our results indicate that the D414V MFN2 variant behaves differently than the few other CMT2A variants that have been investigated in the context of lipid droplets. Furthermore, in contrast to complete loss of MFN2 function, which seems to increase lipid accumulation, the reduced lipid droplet abundance we found in D414V fibroblasts could be due to reduced lipid storage or increased lipid consumption. We suggest that this reduced lipid storage is most likely due to impaired lipid droplet tethering, given the reduced oxidative phosphorylation we observed is inconsistent with an increase in lipid consumption. Furthermore, the distinct distribution pattern of lipid droplets in D414V fibroblasts is consistent with a role of MFN2 regulating interactions between mitochondria and lipid droplets via perilipin (34), an important mediator of lipid droplet distribution (84). In contrast, the other CMT2A MFN2 variants investigated previously are proposed to increase lipid droplet signal via alterations to MERCs (37). Though we also see evidence of MERC disruption in D414V cells, the differences in lipid droplet signal between the D414V and other CMT2A variants could also be due to the severity of MERC disruption, which cannot be compared across studies, as different methods of analysis were used.

In comparison to the characteristics of previous published CMT2A variants, our findings describing the cellular characteristics of the D414V variant, begin to provide insight that may explain the different patient phenotypes and the underlying mechanisms of disease. Though mounting evidence clearly shows that dysregulation of MFN2 causes CMT2A, the exact molecular mechanism underlying this pathology is complicated by the fact that MFN2 is a multifunctional protein. As such, impairment of any or all functions performed by MFN2 could be pathogenic. Here, our description of the D414V variant suggests that impairment of different MFN2 functions may be associated with different pathological phenotypes. Notably, the patient phenotypes described here with the *MFN2* D414V variant are reminiscent of variants in *OPA1*, where patients also have optic atrophy, hearing loss and ataxia. This similarity suggests that impaired mitochondrial fusion may be the underlying mechanism driving these specific phenotypes.

To date, only a few CMT2A MFN2 variants have been investigated for their effects on the various functions of MFN2, making it difficult to generalize which MFN2 function(s) may be causative for which phenotype. Nonetheless, the conflicting findings of pathogenic CMT2A MFN2 variants that have been studied with respect to their consequences on mitochondrial fusion, mtDNA depletion, and bioenergetics suggest that these functions may not be the primary mechanisms underlying the peripheral neuropathy phenotype. However, impairment of these other functions could certainly contribute to disease and may explain some of the additional features sometimes associated with CMT2A (e.g. hearing loss, optic atrophy or lipomatosis). It is also important to note that many of these conflicting reports are from distinct studies by different groups that do not always use the same methods, making direct comparisons difficult.

In summary, our study establishes that *MFN2*-D414V variant is incompetent in carrying multiple *MFN2* functions, arguing for the importance of the HR1 domain. One of the key cellular differences between D414V and CMT2A variants pertains to lipid droplets abundance, as well as a previously unreported perinuclear lipid droplet distribution. Furthermore, the fact that nearby variants also cause distinct phenotypes compared to ‘classic’ CMT2A variants when modelled in flies supports the notion that alterations to the HR1 domain could have different patient phenotypes in humans. However, as only a single D414V patient has been described to date, we cannot infer definitively on genotype/phenotype correlation. At the same time, the cellular phenotypes we describe are all consistent with impaired function of MFN2. Combined with bioinformatic predictions that this variant is likely to be deleterious, we believe the homozygous D414V variant to be the cause of the patient phenotypes, thus expanding the clinical spectrum *MFN2*-associated mitochondrial diseases.

## Materials and Methods

### Case report

The case was identified from the clinical practise of one of the authors (GP). The chart and clinical investigations were reviewed retrospectively to produce the summary provided in this report. The patient provided written informed consent for participation in research, for exome sequencing, and for a skin biopsy in order to isolate primary fibroblast cells which were used in this study. All research was part of studies approved by the University of Calgary Conjoint Health Research Ethics Board.

### Exome sequencing and bioinformatics

DNA was extracted from blood collected into EDTA tubes using standard protocols. Library preparation proceeded using the Ion Ampliseq Exome RDY Panel (Thermo Fisher) according to manufacturer’s protocol. Automated chip loading and templating used the Ion Chef system and 540 chip/chef kit, and sequencing was performed on an Ion S5 system (Thermo Fisher), according to manufacturer’s protocols. Base calling, read alignment to hg19, coverage analysis, and variant calling were performed with Torrent Suite (v. 5.10.1; Thermo Fisher)(85). Patient VCFs were annotated for predicted variant consequence, gnomAD allele frequency (86), CADD score (87), and OMIM phenotypes (88), in addition to default parameters with Ensembl’s command line Variant Effect Predictor (VEP) (89).

### Cell maintenance

Control and patient fibroblast cultures were generated from skin biopsies and cultured in MEM media (Gibco, 11095080) containing l-Glutamine and supplemented with 10% fetal bovine serum (FBS) and 1 mM sodium pyruvate. Cells were maintained at 37°C and 5% CO2.

### Immunofluorescence staining and microscopy

Fibroblasts were seeded on glass coverslips (Fisherbrand, 12-545-81) placed in 24-well plate, at a density of 2 × 10^4^ cells per well, and incubated for 1–2 days. Subsequently, cells were washed with 1xPBS (37°C) and fixed with 4% paraformaldehyde (37°C), permeabilized with 0.1%TritonX-100, blocked with 5% FBS and then the target proteins were probed with primary and appropriate fluorophore conjugated secondary antibodies (Thermo Fisher Scientific), as previously described (Sabouny et al., 2019). Phosphate buffered saline (PBS) was used to wash cells between the steps, and to prepare all reagent solutions. Primary antibodies used are, anti-TOMM20 (Santa Cruz Biotechnology, F-10), anti-dsDNA (autoanti-dsDNA, deposited by Voss. E.W. at DSHB). Immunostained cells mounted on glass slide, and imaged on an Olympus spinning disc confocal system (Olympus SD OSR) (UAPON 100XOTIRF/1.49 oil objective) operated by Metamorph software. Z-stacks of cells were acquired and their z-projection images were used for data analysis.

### Mitochondria-ER contact sites analysis by proximity ligation assay (PLA)

Number and size of MERCs was analyzed using the kit-‘Duolink^®^ In Situ Proximity Ligation Assay’, as described previously (Tubbs and Rieusset, 2016). Briefly, fibroblasts cultured on glass coverslips were washed, fixed and permeabilized. Blocking of unspecific binding sites was performed (1 h at 37°C in a humidified chamber). Then the primary antibodies, TOMM20 (rabbit, Sigma-Aldrich HPA011562-100U) and Calnexin (mouse, EMD Millipore, MAB3126) were applied at 1:1000 dilution. Then the cells were incubated with oligonucleotide conjugated secondary antibodies (anti-rabbit PLUS #DUO92002, and anti-mouse MINUS #DUO92004) diluted 1:5 for 1 h at 37°C in a humidified chamber), followed by ligation and amplification steps using the detection reagents red kit (DUO92008). Thereafter, the primary antibodies were further labelled with appropriate Alexa Fluor conjugated secondary antibodies. Coverglasses with immuno-stained cells were mounted on a glass slide with ProLong™ Glass Antifade Mountant with NucBlue™ Stain (Thermo Fisher Scientific, P36983). Z-stack were acquired with multichannel confocal microscopy for signals from PLA, DAPI and labelled TOMM20 as well as calnexin. Maximum intensity projections of the Z-stacks containing the PLA signals were analyzed with ImageJ FIJI using its ‘Analyze particle’ tool to retrieve the number and sizes of the MERCs.

### Lipid droplets staining

Fibroblasts grown on glass coverslips were fixed with 4% PFA for 20 min at 37°C. After washing off the PFA, cells were permeabilized using 0.1% Saponin (Sigma Aldrich, SAE0073-10G) for 15 min at 37°C. Then the neutral lipid dye, HCS LipidTox green (Thermo Fisher Scientific, # H34350), was applied at a dilution of 1:1000 and incubated overnight at 4°C. The dye solution was washed off and coverslips were then blocked 10% FBS in PBS. Stained cells were washed with PBS and mounted on glass slides with ProLong™ Glass Antifade Mountant with NucBlue™ Stain (Thermo Fisher Scientific, P36983), and microscopy was performed as aforementioned.

### Image Analysis

Mitochondrial network morphology was quantitatively analyzed by measuring the mitochondrial lengths using ImageJ FIJI. Briefly, background was subtracted, then images of mitochondrial networks were skeletonized and the analyze skeleton function was used to obtain the mitochondrial length. The sum of the lengths of all branches of a mitochondrion was evaluated as the total length of that mitochondrion. For each of control or patient fibroblast, at least 20 cells were evaluated. The results shown are one of three independent biological replicates with the same trends showing mean ± SD, and P values based on unpaired, 2-tailed Student’s t-tests.

Size and number of mitochondrial nucleoids as well as MERCs were analyzed using the ‘Analyze particle’ tool in ImageJ FIJI (90). Mitochondrial nucleoids or PLA signals (indicating MERCs) in the fibroblasts were evaluated from the maximum intensity projection of the z-stacks by using ‘Analyze particle’ function of ImageJ FIJI (nuclear signal was excluded from images of fibroblasts immunostained with anti-DNA antibody). The analyses were performed on at least 20 fibroblasts for each patient and control lines. The results represent mean ± SD, and P values were based on unpaired, 2-tailed Student’s t-tests. Each data point is presented as the number of nucleoids per cell or the average size of all nucleoids per cell. In most cases there was one cell per image, but in some cases there were multiple cells per image. In the latter case the numbers were averaged for the number of cells in the image. Bar graphs indicate the average nucleoid sizes or counts in all the fibroblasts analyzed ± SD. P values were based on unpaired, 2-tailed Student’s t-tests.

Lipid droplet numbers were calculated using the same procedure for analyzing the numbers of mitochondrial nucleoids and PLA signals as described above. The distance of individual lipid droplets from nucleus was calculated using ImageJ FIJI. Briefly, the z-projection (maximum intensity) images of cells stained with the LipidTox Green dye and DAPI were processed with the Analyze Particle function to get coordinates for the centre of the individual lipid droplets, as well as the nucleus. These co-ordinates were used to calculate the distance between the centre of each lipid droplet and the center of the nucleus. The distance of each lipid droplet from the nucleus was calculated for at least 20 cells, and used to determine either the distance distribution for all lipid droplets, or the average lipid droplet distance per cell.

### mtDNA copy number analysis

Total DNA (nuclear and mitochondrial DNA) was purified from control and patient fibroblasts (seeded at 5 × 10^5^ cells) using the PureLink Genomic DNA Mini Kit (Thermo Fisher Scientific, K182001) according to manufacturer’s instructions. The relative mtDNA copy number was assessed using QuantStudio 6 Flex Real-Time PCR system (Thermo Fisher Scientific). The mtDNA and the nuclear-encoded housekeeping gene 18S were amplified using primer sequences, and thermocycling conditions exactly as described in (91). Briefly, the 20 μL quantitative PCR (qPCR) reaction contained 10 μL PowerUp SYBR Green Master Mix (Thermo Fisher Scientific, A25742), 100 ng total DNA as template and 500 nM forward and 500 nM reverse primers (final concentrations). MtDNA copy number relative to 18S was analyzed using the delta delta Ct method and represented as percent control (92). Reactions were performed in triplicate technical replicates. Data is presented as mean ± SD and unpaired, 2-tailed Student’s t-tests were used to determine statistical significance.

### Long range PCR

Long range PCR reactions were performed to examine mtDNA deletions as reported previously (Nishigaki et al., 2004). The following primers were used to amplify nearly full length mtDNA (16.3 kb), (1482–1516 F: ACCGCCCGTCACCCTCCTCAAGTATACTTCAAAGG; 1180–1146 R: ACCGCCAGGTCCTTTGAGTTTTAAGCTGTGGCTCG).

The Takara LA Taq polymerase (Takara Bio, RR002M) was used with 250 ng genomic DNA and 200 nM forward and reverse primers. Cycling conditions for the PCR were as follows: 94°C for 1 min; 98°C for 10 s and 68°C for 11 min (30 cycles); and a final extension cycle at 72°C for 10 min. PCR products were visualized by electrophoresis on a 0·6% agarose gel, run for approximately 12 h at 20 V.

### Mitochondrial respiration

Mitochondrial oxygen consumption rates (OCR) in control and patient fibroblasts were measured using a Seahorse XFe24 Extracellular Flux Analyzer (Agilent Technologies, Inc) as described previously (21). Briefly, cells were seeded in an XF24 microplate (3.75 × 10^4^/well) and incubated at 37°C, 5% CO2 for 24 h. Prior to measurement, the growth media was replaced and cells equilibrated in assay media supplemented with d-Glucose (25 mM), sodium pyruvate (2 mM) and l-Glutamine (4 mM). Oxygen consumption rates were calculated following injection of the following compounds: oligomycin (1 μg/mL) (Enzo Life Sciences, BML-CM111), carbonyl cyanide 4-(trifluoromethoxy) phenylhydrazone (FCCP, 1 μM) (Enzo life Sciences, BML-CM120) and Antimycin A (1 μM) (Sigma Aldrich, A8674). Data were normalized to protein content data for each well measured by BCA assay (Thermo Fisher Scientific, 23225).

## Acknowledgements

We would like to thank the patient and their family for participation.

## Author Contributions

Conceptualization: G.S., R.S., G.P., T.S.; Methodology: G.S., R.S., K.M.; Formal analysis: K.M. M.J., A. dK., G.P.; Investigation: G.S., R.S. M.J.; Resources: D.M., G.P.; Data Curation: M.J.; Writing – original draft: G.S., G.P., T.S.; Writing – review & editing: G.S., R.S., D.M., G.P., T.S.; Visualization: G.S.; Supervision: A. dK., G.P., T.S.; Project Administration: G.P., T.S.; Funding acquisition: T.S.

## Abbreviations

HR1: Heptad Repeat 1
HR2: Heptad Repeat 2
MFN: Mitofusin
CMT2A: Charcot-Marie Tooth type 2A
OPA1: Optic Atrophy 1
IMS: Inter Membrane Space
mtDNA: Mitochondrial Dexoyribo Nucleic Acid
ER: Endoplasmic Reticulum
MERCs: Mitochondria – Endoplasmic Reticulum Contact sites
CAPOS: Cerebellar ataxia, Areflexia, Pes cavus, Optic atrophy, and Sensorineural hearing loss
GAD: Glutamic Acid Decarboxylase
MRI: Magnetic Resonance Imaging
NGS: Next Generation Sequencing
PLA: Proximity Ligation Assay
BDLP: Bacterial Dynamin Like Protein
OCR: Oxygen Consumption Rate
EDTA: Ethylenediaminetetraacetic acid
VCFs: Velocardiofacial syndrome
CADD: Combined Annotation Dependent Depletion
OMIM: Online Mendelian Inheritance in Man
VEP: Variant Effect Predictor
CO2: Carbon dioxide
MEM: Minimum Essential Media
FBS: Fetal Bovine Serum
PBS: Phosphate Buffered Saline
DSHB: Developmental Studies Hybridoma Bank
DAPI: 4′,6-diamidino-2-phenylindole
PFA: Paraformaldehyde
SD: Standard Deviation
qPCR: Quantitative Polymerase Chain Reaction

